# Carotenoid-dependent plumage coloration is associated with reduced male care in passerine birds

**DOI:** 10.1101/2022.12.01.518672

**Authors:** Verónica A. Rincón-Rubio, Tamás Székely, András Liker, Alejandro Gonzalez-Voyer

**Affiliations:** Departamento de Ecología Evolutiva, Instituto de Ecología, Universidad Nacional Autónoma de México, México, Mexico; Posgrado en Ciencias Biológicas, Universidad Nacional Autónoma de México, México, Mexico; Milner Centre for Evolution, Department of Biology and Biochemistry, University of Bath, Bath, BA2 7AY, UK; Department of Evolutionary Zoology, University of Debrecen, Debrecen, H-4032, Hungary; ELKH-PE Evolutionary Ecology Research Group, University of Pannonia, Veszprém, Hungary; Behavioural Ecology Research Group, Center for Natural Sciences, University of Pannonia, Veszprém, Hungary

**Keywords:** Carotenoid-dependent coloration, sexual selection, parental care, passerines, evolution, animal signals

## Abstract

The immense diversity of plumage coloration exhibited by birds is the result of either pigments deposited in the feathers or microstructural arrangements of feather barbules. Some of the most common pigments are carotenoids that produce bright yellow, orange and red colors. Carotenoids differ from other pigments since birds cannot synthesize them de novo and must obtain them from the diet. Carotenoid pigments are usually associated with signaling and sexual selection, although they also have antioxidant properties and play a role in the immune response. Here we hypothesize that carotenoid-dependent plumage coloration functions as a signal of a male’s tendency to invest in offspring care because they play an important role in the self-maintenance and may provide key information about individual quality; allowing females to obtain information about a males’ tendency to invest in offspring care. Using phylogenetic comparative analyses across 350 passerine birds we show that species that consume carotenoid-rich foods have more carotenoid-dependent plumage coloration than species with carotenoid-poor diets. In addition, carotenoid-dependent plumage coloration is associated with a decreased male investment in offspring care. Our results suggest investment into carotenoid-dependent plumage coloration trades off against male investment into offspring care and will likely have broad implications for our understanding of the ecological contexts that facilitate various evolutionary processes such as sexual selection or signaling associated with plumage colors.

## INTRODUCTION

Animals employ a diversity of signals to transmit information about quality and receptivity to potential mates, and birds are no exception. Birds have a diverse repertoire of sexual signals, ranging from simple to rather intricate courtship dances, to elaborate songs and colorful ornaments such as plumage coloration. These diverse signals vary in their costs of production, which can determine their function (Badyaev and Hill 2000; Senar et al. 2002). Sexual signals are useful to receivers if the information they transmit is honest. Honest-advertisement models of sexual selection predict that low-quality individuals are precluded from signaling at high levels because sexual signals are costly to produce and maintain, and because such costs are prohibitively larger for low-quality versus high-quality individuals (Zahavi 1975, 1977; Anderson 1994). Sexual signals used in mate choice are predicted to be under directional selection, favoring increasingly more exaggerated signals, due to intra-sexual competition or mate preferences, with survival costs imposing a cap on the degree of elaboration (Anderson 1994). The elevated costs and directional selection acting on sexual signals seem to run counter to the observation that many animals present multiple signals (e.g. song and plumage coloration in birds), as high costs would be expected to limit investment into more than one signal (Iwasa and Pomiankowski 1994; but see Gonzalez-Voyer et al. 2013). Theory suggests that multiple signals could evolve if they reflect a different aspect of the overall condition of the emitter (Møller and Pomiankowski 1993). Nonetheless, the ultimate reasons why birds use more than one single signal remain difficult to explain and the striking diversity of signals in nature begs the question: What are all these signals communicating?

Many tests of the signaling function of ornamental traits have focused on visual signals, and in birds in particular colorful feathers (e.g. Butcher and Rohwer 1989; Savalli 1995). Plumage coloration is the most conspicuous secondary sexual ornament of many bird species (Toews et al. 2017). Among the mechanisms responsible for plumage coloration are pigment-based ornamentation, and the most prevalent pigments in the avian integument are melanins, which result in mainly dark and brownish colors, and carotenoids, which usually result in bright yellow, orange or red colorations (Hill & McGraw 2006). The latter must ultimately be obtained from the diet as animals cannot synthesize carotenoids de novo (Olson and Owens 2005; Weaver et al. 2018). Recent work with the Tyrannida suboscine radiation found that rapid rates of plumage color divergence in males were biased towards carotenoid-dependent colors, in line with their assumed role in sexual signaling (Cooney et al. 2019).

Carotenoids are not only used as pigments but also play an important role in the self-maintenance of individuals, as they are involved in numerous physiological pathways, such as immunocompetence, and vitamin A synthesis, and play a role as antioxidants (Latscha 1990; Weaver et al. 2018). The ability to use carotenoids present in the diet may be limited by physiological (Negro et al. 1998; Saino et al. 1999), and nutritional constraints (Hill 2000), as well as by environmental agents such as intestinal parasites interfering with carotenoid absorption (McGraw and Hill 2000). Due to the diversity of functions of carotenoids in avian physiology, including but not limited to combating oxidative stress, many of the classic examples of honest signaling in animals are based on carotenoid-dependent coloration (e.g. Hill 1991; Burley et al. 1992; Metz and Weatherhead 1992). Indeed, researchers hypothesized that assessment of the carotenoid-dependent coloration of prospective mates provides key information about individual quality and enables choices that increase fitness (McGraw 2006; Weaver et al. 2018).

The benefits associated with mate choice involve both indirect benefits, e.g. good genes for offspring, or direct benefits, for example when both mates participate in rearing offspring (Mitchell et al. 2007). Post-hatching parental care is common in birds and evolved to increase offspring survival, although it usually involves a cost to the caregiving parent (reviewed in Vági et al. 2019; Clutton-Brock 2019). In species where both parents invest in offspring care (biparental care), a potentially important benefit for one parent is the level of investment from their partner, which reduces the overall cost of parental care (Trivers 1972; Thornhill 1976; Mitchell et al, 2007), although conflict between mates regarding the amount of investment also occurs (Houston et al. 2005; Osorno and Székely 2004). Because parental performance cannot usually be observed before mate choice, the condition of the prospective mate must be appraised using indirect sexual traits that honestly indicate phenotypic or genetic quality (Kodric-Brown and Brown 1984). However, intraspecific studies in birds offer contrasting evidence regarding whether coloration is a useful indicator of a mate’s investment in parental care (Williams 1966; Hoelzer 1989; Burley 1986; Hill 1991; Magrath and Komdeur 2003). Thus, it is unclear whether carotenoid-dependent coloration in bird plumage may function as a signal of individual quality and in particular direct benefits in terms of investment into parental care by mates, which could shed some light on the immense variation of plumage coloration across birds. We hypothesize that evolution will favor honest signals (e.g., by carotenoid-dependent coloration) in species with biparental care that allow females to obtain information about how liable males are to invest in their future offspring. Carotenoid-dependent plumage coloration could be an indicator of a male’s foraging ability and condition, given they can obtain enough of these phytonutrients to use them in advertisement. Furthermore, a colorful male is possibly more likely to bring carotenoid-rich foods to offspring. We, therefore, predict that carotenoid-dependent plumage coloration will be both present and more abundant in bird species where males actively participate in parental care compared to those in which males do not participate.

We tested this hypothesis using 350 passerine species. Passeriformes is the most recent, diverse, and widely distributed order of birds (Jetz et al. 2012). Thomas et al. (2014) estimated that at least 41% of passerine species show carotenoid-dependent coloration in their plumage. Passerines species extract and accumulate carotenoids more efficiently compared to other groups of birds (McGraw 2005). We assessed the relative contribution of males in parental care and the presence and extent of carotenoid-dependent coloration in the plumage of passerine species while controlling for phylogenetic relationships. We also analyzed their diet, body size, geographical distribution, and polygyny because these could potentially confound the relationship between coloration and male care. Based on recent results by Cooney et al. (2022), we expect a negative relation between carotenoid-dependent coloration and body size and a positive relation with a tropical distribution. We also contemplate the mating system because the social mating system affects male care (o vice versa) hence can be a confounding variable on parental care and the expression of a sexual signal.

## MATERIALS AND METHODS

Data were collected from published sources, including both primary reference works (e.g. Olson and Owens 2005; Thomas et al. 2014; Liker et al. 2015) and compendia such as the Handbook of the Birds of the World Alive (del Hoyo et al. 2016; accessed between May and December 2016). We initially limited our dataset to the 792 species for which Liker et al. (2015) provide detailed information on the relative investment by each sex in parental care. We selected information on relative investment in parental care duties by males of all passerine species for which data was available and combined it with available information on the presence of / amount of carotenoid-dependent coloration in feathers, diet, body size, geographical distribution, and mating system. The final dataset included 350 species and represented a significant proportion of passerine taxonomic diversity (66 families). Information on parental behavior, carotenoid-dependent coloration, and diet were available for almost all species, although sample sizes vary for other traits (see below) due to data availability.

### Parental care

Liker et al. (2015) compiled detailed information on relative investment by each sex in six components of avian parental care from reference books, and published literature (references in Liker et al. 2015 updated in Gonzalez-Voyer et al. 2022). We focus here on only three components: nest building, incubation, and chick feeding, which are the ones for which information was available for the greatest number of species. Nonetheless, mean scores calculated from our set of three care components were highly correlated with the mean scores of the set of six care components (r= 0.8, p= 0.001, N= 390). For each care component, relative participation by males was scored by Liker et al. (2015) on a 5-point scale: 0: no male care, 1: 1–33% male care, 2: 34–66% male care, 3: 67–99% male care, 4: 100% male care. These scores were based on quantitative data, when available (e.g. percentage of incubation by males), or on qualitative descriptions of care in species where quantitative data were lacking. For each species, we calculated a mean relative male participation from the three components.

### Carotenoid-dependent coloration

We used an existing dataset (Thomas et al. 2014) on the presence of carotenoid-dependent plumage coloration in all neornithine species. Presence (1) or absence (0) of carotenoid-dependent coloration was scored for all species based on plumage colors observed in illustrations taken from the Handbook of the Birds of the World Alive online (del Hoyo et al. 2016), using as a reference a set of species for which the presence of carotenoids in plumage had been chemically confirmed. We selected all passerine species from the list and ensured that it was the male who presented carotenoid-dependent coloration. As we are not only interested in the presence of carotenoid-dependent coloration but also in the amount expressed on the body of the species, we developed our own scoring system following Owens and Hartley (1998) to obtain a quantitative estimate for the amount of area of the body with plumage with carotenoid-dependent coloration. For species for which Thomas et al. (2014) had confirmed the presence of carotenoid coloration we estimated the amount of carotenoid-dependent coloration, whereas for those species without carotenoid-dependent coloration according to Thomas et al. (2014) we assigned a value of 0. We estimated the amount of carotenoids distributed throughout the body of the bird because the more pigments you need, the more expensive the signal is likely to be in these species, and we assume the selection pressure will be stronger as well. We only took into account the colorations that typically depend on these pigments: bright reds, oranges, and yellows. To minimize bias, we selected three volunteers with birdwatching experience, but uninformed about the hypotheses we wished to test, to perform the scoring. Each observer was shown illustrations from the Handbook of the Birds of the World of the 177 passerine species that Thomas et al. (2014) classified as presenting carotenoid-dependent plumage coloration and asked to score patches with 1: if the carotenoid-dependent coloration covered the whole patch or a part of it; 0: if it did not show carotenoid-dependent coloration. The observers scored eight body regions per species: head, nape, back, rump, throat, chest, belly, tail, and wings. In the rare instances where the patch observers were meant to score was not visible in the images, we searched descriptions and pictures of the species to verify whether they present carotenoid-dependent coloration in said patch. We scored these regions because they cover the whole body of the bird. In cases where multiple subspecies were illustrated, we scored coloration in the nominate subspecies. The continuous measure for carotenoid-dependent coloration was the sum of the scores for all patches.

Scoring coloration with handbook plates represents a valid alternative to measuring carotenoid color (Dale et al. 2015). Because illustrations are a suitable reference for field identification, special care is taken to reproduce coloration and patterning accurately. Several previous comparative studies have used coloration in plates to test hypotheses about avian color evolution (Olson and Owens 2005; Thomas et al. 2014), and previous works have found that coloration in plates is highly correlated with coloration in museum specimens (Dale et al. 2015).

### Co-variates

To assess the carotenoid content in the diet of each species, we developed a scoring system following Olson and Owens (2005), who scored carotenoid intake based on a coarse-scale index ranking seven diet categories based on their carotenoid content: 1 for seeds, nuts, wood; 2 for nectar, pollen, sap, exudates, lerps; 3 for vertebrates; 4 for invertebrates; 5 for foliage, flowers, fungi; 6 for fruit; and 7 for algae, diatoms. We collected diet data from verbal descriptions in the Handbook of the Birds of the World Alive (del Hoyo et al. 2016). Because most species do not feed solely on a single group, we assigned relative values according to the importance of each group in their diet based on the HBW descriptions. We multiplied the relative values for each group by the amount of carotenoids contained and then added them to obtain the total carotenoids consumed in 100g of food for each species. We then calculated a mean score of carotenoid content from the relative importance of each diet category multiplied by the carotenoid content in that category (from Olson and Owens 2005). We summed the values across all diet categories, for each species. For example, *Malurus coronatus* feeds only on insects, so the amount of carotenoids in the insect group is directly multiplied by 1, giving a total of 0.007g/100g of carotenoids. In contrast, *Piranga olivacea* feeds mainly on insects but occasionally eats small fruits, so we multiplied the amount of carotenoids in the insect group by 0.8 and the amount of carotenoids in the fruit group by 0.2, to finally add the result of both multiplications. The value of 0.8 was maintained in all foods that the literature specifies as main in the diet, while foods described as “sporadic” were given a value of 0.2; values that add up to 1 according to the importance of the group in their diet. In the cases where there was more than one main or sporadic group, 0.8 and 0.2 were divided, respectively, among the total number of food groups ingested.

We used body mass as an estimate of the body size of the species. We included body size in our analyses because it is strongly correlated with life history traits and could reflect possible restrictions for the development of plumage coloration in birds, such as metabolic rate and the ingestion of food with carotenoids. Body mass data were obtained from Liker et al. (2015) for almost all species. For species lacking data, we extracted it from the HBW. Log-transformed values of body mass were used in the statistical analysis.

We obtained the species’ breeding distribution from del Hoyo et al. (2016) for latitude. Geographical distribution has been suggested to potentially influence plumage coloration because of diverse mechanisms, which differ between tropical and non-tropical zones (e.g., parasitic load, and diversity of foods with carotenoids; Møller 1998; Cooney et al. 2022). Because we were mainly interested in the contrast between tropical and non-tropical areas, and geographical distribution was included as a potential confounding factor, species were categorized based on whether they presented a tropical distribution (A: 23° N to 23° S), a non-tropical distribution (B: north of 23° N, or south of 23° S), or the distribution spanned both tropical and non-tropical areas (C: for species breeding in non-tropical and tropical areas).

Because the variance in mating success, a more precise estimate of the intensity of sexual selection, has not been reported for a broad range of species, we use polygamy frequency as a proxy for variance in mating success, assuming that more frequent polygamy by males means higher variance in mating success. Polygamy frequency data for males was obtained from Liker et al. (2015) who scored the overall incidence of social polygamy for each sex on a scale from 0 to 4, with 0 corresponding to no (or very rare) polygamy (< 0.1% of individuals), 1 to rare polygamy (0.1-1%), 2 to uncommon polygamy (1-5%), 3 to moderate polygamy (5-20%), and 4 to common polygamy (>20%). These scores explain a high proportion of the variation in the actual polygamy frequencies (Liker et al. 2015).

### Comparative analyses

To account for phylogenetic relationships among passerine birds for all analyses we created a Maximum clade credibility tree from 1000 phylogenetic trees randomly selected from 10,000 phylogenies from the pseudo-posterior distribution from the avian phylogeny of Jetz et al. (2012; available at http://birdtree.org). To create this consensus tree, we used the package *phangorn* version 2.10.0 (Schliep et al. 2011).

Here, we aimed to address two questions. First, we wanted to know if the presence of carotenoid-dependent coloration in feathers, irrespective of the area of the body it covered, serves as an indicator of males’ tendency to invest in parental care. Second, we wanted to assess if the magnitude of the signal (proportion of the body area) could be a more reliable indicator of the males’ tendency to invest in parental care. We, therefore, developed two phylogenetically controlled models. To test only whether males of a given species express carotenoid-dependent coloration, the response variable was binary: presence (1) or absence (0) of plumage carotenoid-dependent coloration. To test the relationship with parental care, and other co-variates, we used a phylogenetic logistic regression for binary dependent variables (Ives and Garland 2010). To test if the signal size (measured as coverage of the body by carotenoid-dependent plumage coloration) is an important indicator of parental care, the response variable was continuously allowing us to use phylogenetic generalized least squares regression (PGLS; Martins and Hansen 1997).

Phylogenetic logistic regression for binary dependent variables estimates the phylogenetic signal through the parameter α that calculates the transition rate in the dependent variable, that is, the speed at which phylogenetic correlations between species are lost. Large values indicate that transitions between 0 and 1 occur quickly, breaking the tendency that closely related species resemble one another, while low values are the opposite (e.g. high phylogenetic signal). These analyses were carried out using the ‘phylolm’ (Ho and Ané 2014) package in R.

In a similar way, PGLS is a linear regression model in which phylogenetic information is incorporated into the error term (residuals; Martins and Hansen 1997). PGLS calculates the measure of phylogenetic correlation in the residuals with the parameter λ (Pagel 1999). λ values vary between 0 and 1, where λ = 0 denotes no phylogenetic signal (independent evolution of traits) and λ = 1 denotes strong phylogenetic dependence (Pagel 1999; Freckleton et al. 2002). These analyses were carried out using the ‘caper’ (Orme et al. 2012) package in R. In all models, we included diet, body size, geographical distribution, and the mating system as co-variates.

## RESULTS

All analysis included 350 of the 5,986 species of passerine birds reported by Jetz et al. (2012). These 350 species represent 5.8% of the named species richness of the order. The species are distributed across 66 families, so the analysis included 66% of the families reported for the order Passeriformes. The biggest limitation to increasing the sample size is the lack of information on male investment in parental care since it is quite detailed and therefore not available for many species.

Both the extent of carotenoid-dependent coloration and paternal care vary between passerines. First, the extent of male care varies among species, from 1 (e.g. *Cincloramphus cruralis)* to 99% (e.g. *Remiz pendulinus*) of total care, although in no species males are the sole care providers. Second, in 45% of the species males exhibit carotenoid-dependent coloration in their plumage (Figure 1).

**Figure 1.**
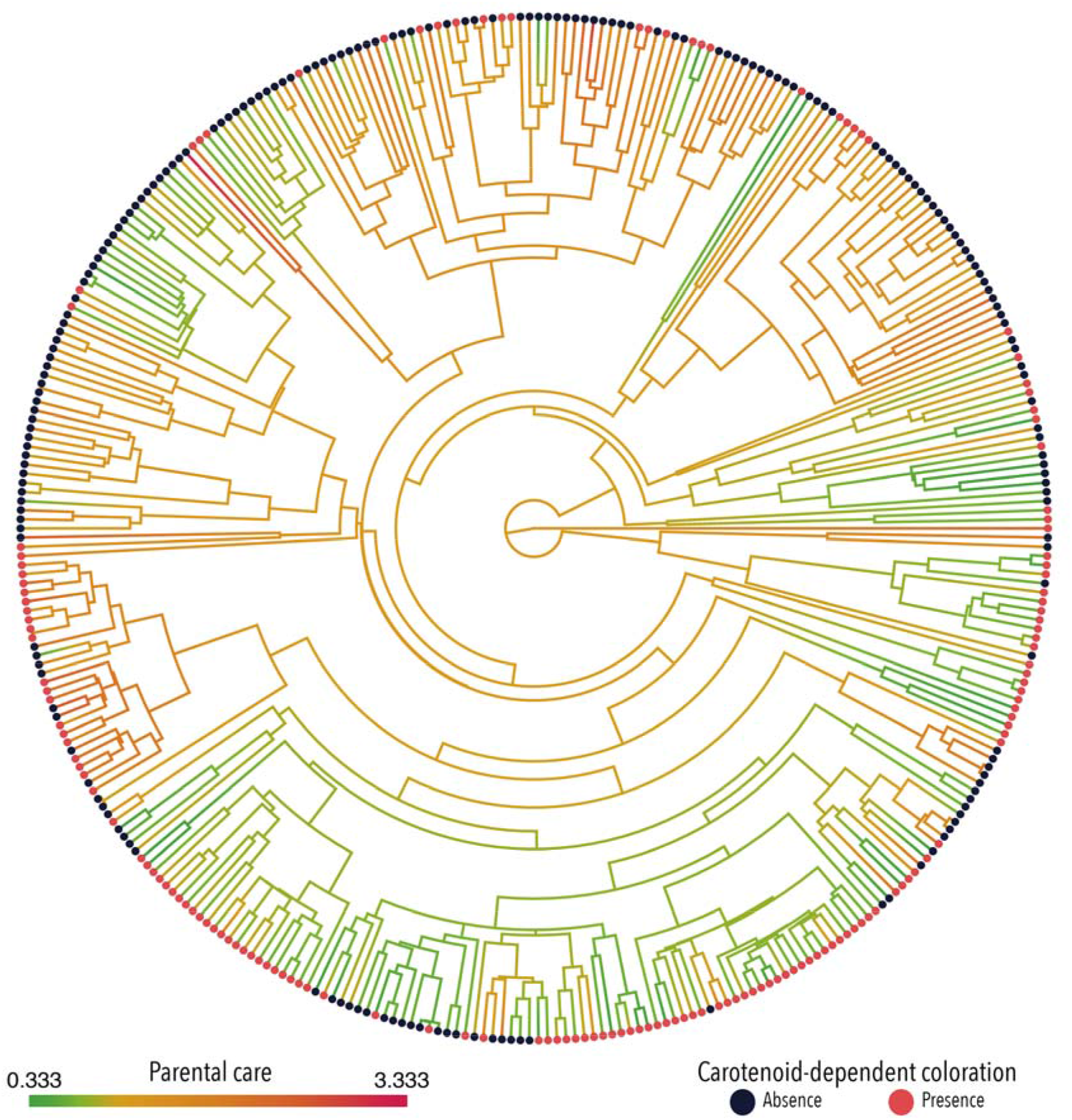
Phylogenetic distribution of male investment into offspring care (reconstructed along the branches of the phylogeny) and presence of carotenoid-dependent coloration (red for presence and blue for absence) in passerine birds (n= 350, species phylogeny from Jetz et al. 2012). Species names are omitted for clarity of the figure.

### Presence of carotenoid-dependent coloration

Males participate significantly less in offspring care when they present carotenoid-dependent coloration in their plumage (Table 1), contrary to our hypothesis. Diet was also significantly positively associated with carotenoid-dependent coloration, as predicted and in line with previous results (Olson and Owens 2005), indicating that species presenting carotenoid-dependent coloration in their plumage have a more carotenoid-rich diet than species who do not present carotenoid coloration in their plumage. Neither polygyny nor geographic distribution predicted carotenoid coloration (Table 1). Although body size was not significantly associated with the presence of carotenoid coloration in the plumage, we cannot reject an association because the confidence interval does not overlap with 0 (Table 1; see Ives and Garland 2010), suggesting that smaller species tend to be more likely to present carotenoid coloration than larger ones.

**Table 1.** Presence of carotenoid-dependent coloration (response variable) in relation to the extent of paternal care and carotenoid content in the diet using phylogenetic logistic regression for binary dependent variables. Body size, polygamy and distribution are included as potential confounding variables. Confidence intervals (CIs) of the slopes were calculated at 95%, n= 350 species, α= 0.02. The table shows the slope and its standard error (Std Err) for each predictor, the *z* statistic, upper and lower confidence intervals and the p value.

| Predictors | Slope | Std Err | $z$ | Lower CI | Upper CI | $p$ |
| --- | --- | --- | --- | --- | --- | --- |
| Intercept | 0.17 | 0.82 | 0.21 | -0.91 | 1.59 | 0.84 |
| Paternal care | -0.41 | 0.20 | -2.03 | -0.80 | -0.06 | 0.04* |
| Diet | 23.81 | 8.01 | 2.97 | 23.64 | 24.06 | 0.003* |
| Body size | -0.53 | 0.34 | -1.54 | -1.03 | -0.10 | 0.12 |
| Polygamy | 0.01 | 0.06 | 0.15 | -0.13 | 0.10 | 0.88 |
| Distribution B | 0.24 | 0.36 | 0.69 | -0.51 | 0.83 | 0.49 |
| Distribution C | 0.13 | 0.33 | 0.39 | -0.61 | 0.66 | 0.70 |

\*Significant at the 0.05 probability level

### The extent of carotenoid coloration

The results of the PGLS generally confirmed those of the logistic regression. We found that the area of the body covered by carotenoid-dependent plumage coloration increases significantly with higher carotenoid content in the diet (Table 2). Furthermore, males who invest less in paternal care present significantly more carotenoid-dependent coloration in their plumage (Table 2). Finally, the proportion of the body area with carotenoid-dependent plumage coloration was unrelated to body size, polygyny, or geographic distribution (Table 2).

**Table 2.** Presence of carotenoid-dependent coloration (response variable as a continuous measure) in relation to the extent of paternal care and carotenoid content in the diet using PGLS. Body size, polygamy and distribution are included as potential confounding variables. n= 350, λ = 0.79. The table shows the slope (β) and its standard error (Std Err), the *t* statistic and p value for each predictor.

| **Predictors** | **$\beta$** | **Std Err** | **$t$** | **$P$** |
| --- | --- | --- | --- | --- |
| Intercept | 0.32 | 0.20 | 1.64 | 0.14 |
| Parental care | -0.07 | 0.03 | -1.96 | 0.05* |
| Diet | 3.31 | 1.22 | 2.72 | 0.01* |
| Body size | -0.09 | 0.06 | -1.50 | 0.13 |
| Polygamy | 0.002 | 0.01 | 0.17 | 0.87 |
| Distribution B | 0.06 | 0.06 | 0.10 | 0.33 |
| Distribution C | 0.06 | 0.06 | 1.12 | 0.27 |

\* Significant at the 0.05 probability level

## DISCUSSION

To our knowledge, this is the first study to test a relationship between parental care and ornaments across a broad range of species. Here we used two means of quantifying carotenoid-dependent plumage coloration and found similar biological patterns. Our results show two significant associations with carotenoid-dependent plumage coloration: firstly a negative association with paternal care, and secondly a positive link with carotenoid content in the diet, as shown in a previous study (Olson and Owens 2005).

### Negative association between parental care and carotenoid-dependent plumage coloration

Although previous studies reported positive associations between parental care and carotenoid-dependent plumage coloration at a within-species level (Senar et al. 2002; Ewen et al. 2008; Pagani-Nuñez and Senar 2014), we found a negative association when comparing among species. Both positive and negative associations between carotenoid-dependent plumage coloration have been reported in studies with single species, for example in northern cardinals carotenoid-dependent plumage brightness is a positive indicator of parental care (Linville et al. 1998); on the contrary, in house finches elaborately ornamented males with carotenoid-dependent coloration avoid costly parental care (Duckworth et al. 2003). The negative association between male investment in offspring care and carotenoid-dependent plumage coloration is in line with what could be predicted based on a trade-off between a presumably costly sexual signal and another energetically demanding activity (Stearns 1992). A high cost of paternal care is supported by the results of a meta-analysis which found that males that invest more into parental care had reduced survival compared with controls, while females showed no reduction in survival as a result of higher parental care (Santos and Nakagawa 2012). In addition, a comparative analysis across 194 bird species found that higher male-male competition for access to females was associated with increased male mortality and that a mortality cost of parental care is only observed in males, after controlling for the effects of intra-sexual competition, but not in females (Liker and Székely 2005). Our results suggest that carotenoid-dependent plumage coloration may be signaling indirect benefits rather than a greater contribution to offspring care (e.g. Olson and Owens 1998; Hill 1999).

### Benefits associated with carotenoid-dependent plumage coloration

Indirect benefits could be the main cause for the preference for carotenoid-dependent coloration over other coloration mechanisms. Therefore, the evolution of these colorations will depend on their ability to act as signals that indicate the genetic benefits that offspring will inherit, such as attractive offspring or good genetic quality. While our results indicate these signals do not reflect a male’s ability to be a good parent, it has been suggested that carotenoid-dependent colors provide information on the general genetic quality or differences in metabolic capacities among conspecific males (Hamilton and Zuk, 1982; Blount, 2004). In addition, if resistance to oxidative stress is hereditary (Kim et al. 2010), females will be receiving indirect benefits by having offspring with greater resistance. Nevertheless, we cannot exclude the possibility that carotenoid-dependent coloration provides information about other types of direct benefit, such as a higher-quality or more productive nesting site, which would enable females to more easily acquire resources for offspring including carotenoid-rich food (Heywood 1989; Hoelzer 1989; Price et al. 1993). Indeed, territoriality has also been correlated with carotenoid-dependent coloration in intraspecific studies, for example, in red collared widowbirds, floaters and residents differ markedly in both the area and redness of their carotenoid badge, being redder in residents; which reveals an effect of carotenoid signal variation in male contest competition, and represents a strong case for “carotenoid status signaling” in a bird species (Andersson et al. 2002).

### Carotenoid-dependent plumage coloration is associated with carotenoid-rich diets

The positive association between carotenoid-dependent plumage coloration and carotenoid-rich diets is in agreement with the large number of single-species studies that have shown that carotenoid-dependent plumage pigmentation is closely linked to the dietary availability of carotenoids (e.g. Hill et al. 2002; McGraw et al. 2003; Navara and Hill 2003; McGraw et al. 2004; for an example in fishes see: Kodric-Brown 1989), and with the comparative study of Cooney et al. (2022). It was of course to be expected that high availability of dietary carotenoids was a necessary pre-requisite for the presence of carotenoid-dependent plumage coloration, as birds cannot synthesize carotenoids de novo. In addition, a previous comparative study found a significant association between diet and plumage coloration across 140 avian families (Olson and Owens 2005). Previous work showed that diet influences intraspecific variability in plasma carotenoid concentrations of individuals (Bortolotti et al. 2000; Negro et al. 2000) and that such variability is reflected in their expression of plumage coloration (Hill et al. 1994; Bortolotti et al. 1996; Saino et al. 1999; Negro et al. 2000). Such differences may not be solely due to availability of carotenoids in the diet but also to interspecific differences in metabolism and individual differences in absorption, transport, and expression of carotenoids in the plumage. In line with this suggestion, Tella et al. (2003) reported that carotenoid concentration in plasma increases as species’ body size decreases since smaller bird species have higher food intake rates per unit of body mass as required by their high metabolic rates. Our results are in line with those reported by Tella et al. (2003), as we found a negative relationship between carotenoid-dependent coloration and body size.

### Tropical passerines do not present more carotenoid-dependent plumage coloration

Tropical birds are often thought to be more colorful than the rest, but we did not find a significant relationship between carotenoid-dependent plumage coloration and a tropical distribution, in line with results from previous studies (Bailey, 1978). Although Wilson and von Neumann (1972) reported an association between tropical distribution and plumage coloration in birds, they included several non-passerine species. When they repeated the analysis with only passerines, the association between plumage coloration and distribution disappeared. Nonetheless, recent studies (Dale et al. 2015; Cooney et al. 2022) present results that suggest that tropical passerine species are more colorful than species that live in non-tropical areas, but in none of the studies the authors distinguish between different coloring mechanisms. This differentiation is not trivial because, according to Bailey (1978), iridescent colors are more common in tropical areas, but this has not been demonstrated for carotenoid-dependent colorations. We recommend taking more precise measures of distribution in future studies.

### Carotenoid-dependent plumage coloration is not associated with higher polygyny

Having found a significantly negative relationship between carotenoid-dependent plumage coloration and parental care, it may be disconcerting not to observe a relationship between carotenoid-dependent plumage coloration and polygyny. The absence of a relationship may be due to the fact that most passerines have a monogamous mating system (Lack 1968), and therefore polygyny frequency may not be an ideal proxy to measure mating opportunities. In addition, a study with a much broader sample of species found rather weak correlations between different sex role components (that is estimates of intra-sexual competition (sex size dimorphism), mate attraction (sexual dichromatism), mating system and parental care), including a weak correlation between mating system and the investment in parental care by both sexes (Gonzalez-Voyer et al. 2022). Such weak correlations between sex role components could explain why we did not find an association between the mating system and the presence of carotenoid-dependent plumage coloration. It is also possible that considering extra-pair copulations, which are very common in monogamous species such as passerines (Dunn et al. 2001), would provide greater insight into the possible function of carotenoid-dependent coloration in reproductive success, as previous studies have suggested sexual dichromatism is related to extra-pair copulation while sexual size dimorphism plays a role in intra-sexual competition for mates (Owens and Hartley 1998; Gonzalez-Voyer et al. 2022). Extra-pair copulation not only contemplates the opportunity of a male to mate again but also is an indicator of the probability of paternity loss, which in turn disincentivizes male investment into parental care (Kvarnemo 2005).

In conclusion, our study reveals that the presence of carotenoid-dependent coloration is negatively correlated with males’ tendency to invest in parental care, indicating that carotenoid-dependent coloration most likely provides information on potential indirect benefits to females. As predicted, we also found a significant association between the amount of carotenoids in the diet and their use for carotenoid-dependent plumage coloration, both as a binary and continuous measure. The association between diet and plumage coloration suggests a possible co-evolutionary relationship between plants and passerine birds which would be worthwhile exploring. Even though our hypothesis was rejected for carotenoid-dependent plumage coloration, it would be worthwhile testing whether there is an association with tegumentary coloration, such as that expressed in the beak and legs (e.g. Montoya and Torres 2015). Tegumentary coloration could be a better sign of the current condition, unlike plumage coloration which reflects the condition of an individual at the time of molting (Duckworth et al. 2003). Dynamic signals require constant investment and are thus more sensitive to environmental conditions (Dale 2000; Bennett et al. 2002; Blount et al. 2003; Faivre et al. 2003).

## ACKNOWLEDGEMENTS

We are grateful for the valuable assistance of S. Larre, T. Nakamura, and R. Santos-Gally to quantify the extent of carotenoid-dependent coloration. Edgar Ávila Luna provided logistical support. This study was funded by a PAPIIT-UNAM research grant (IN211919) to AG-V and by a Royal Society Newton Advanced Fellowship (NA150257) to AG-V hosted by TS. This work was initiated during Verónica Rincón Rubio’s undergraduate thesis and continued for her Ph.D. in the Posgrado en Ciencias Biológicas, Universidad Nacional Autónoma de México. VRR received a Ph.D. fellowship from CONACyT (924964).

## Notes

### Competing Interest Statement

The authors have declared no competing interest.

